# Adult sex change leads to extensive forebrain reorganization in clownfish

**DOI:** 10.1101/2024.01.29.577753

**Authors:** Coltan G. Parker, George W. Gruenhagen, Brianna E. Hegarty, Abigail R. Histed, Jeffrey T. Streelman, Justin S. Rhodes, Zachary V. Johnson

**Affiliations:** Neuroscience Program, University of Illinois, Urbana-Champaign, Illinois, USA; School of Biological Sciences, Georgia Institute of Technology, Atlanta, Georgia, USA; Department of Psychology, University of Illinois, Urbana-Champaign, Illinois, USA; Department of Psychiatry and Behavioral Sciences, Emory University, Atlanta, GA, USA; Emory National Primate Research Center, Emory University, Atlanta, GA, USA

**Keywords:** Anemonefish, preoptic area, telencephalon, sex difference, sex change

## Abstract

Sexual differentiation of the brain occurs in all major vertebrate lineages but is not well understood at a molecular and cellular level. Unlike most vertebrates, sex-changing fishes have the remarkable ability to change reproductive sex during adulthood in response to social stimuli, offering a unique opportunity to understand mechanisms by which the nervous system can initiate and coordinate sexual differentiation. This study explores sexual differentiation of the forebrain using single nucleus RNA-sequencing in the anemonefish *Amphiprion ocellaris*, producing the first cellular atlas of a sex-changing brain. We uncover extensive sex differences in cell type-specific gene expression, relative proportions of cells, baseline neuronal excitation, and predicted inter-neuronal communication. Additionally, we identify the cholecystokinin, galanin, and estrogen systems as central molecular axes of sexual differentiation. Supported by these findings, we propose a model of neurosexual differentiation in the conserved vertebrate social decision-making network spanning multiple subtypes of neurons and glia, including neuronal subpopulations within the preoptic area that are positioned to regulate gonadal differentiation. This work deepens our understanding of sexual differentiation in the vertebrate brain and defines a rich suite of molecular and cellular pathways that differentiate during adult sex change in anemonefish.

**Significance Statement:** This study provides key insights into brain sex differences in sex-changing anemonefish (*Amphiprion ocellaris*), a species that changes sex in adulthood in response to the social environment. Using single nucleus RNA-sequencing, the study provides the first brain cellular atlas showing sex differences in two crucial reproductive areas: the preoptic area and telencephalon. The research identifies notable sex-differences in cell-type proportions and gene expression, particularly in radial glia and glutamatergic neurons that co-express the neuropeptide cholecystokinin. It also highlights differences in preoptic area neurons likely involved in gonadal regulation. This work deepens our understanding of sexual differentiation of the brain in vertebrates, especially those capable of adult sex change, and illuminates key molecular and cellular beginning and endpoints of the process.

## Introduction

Sex is associated with differences in physiology and behavior across taxa, and in humans is associated with differential risk for many reproductive, endocrine, and neuropsychiatric diseases. Sexual biology is inextricably controlled by the brain, specifically neurons in the preoptic area (POA) of the hypothalamus and interacting forebrain regions that regulate pituitary release of the gonadotropins that in turn regulate the gonads (the HPG axis). Much of our understanding of sex differences in the brain comes from a few mammalian and avian systems in which sexual differentiation is genetically determined, irreversible, and accomplished during early development (1–4). These systems capture only a narrow degree of mechanistic diversity in sexual biology that exists in nature. Research in non-model systems that represent the true diversity of sexual biology found in nature is necessary to define the flexibility and rigidity of specific control mechanisms within the POA and associated regions of the forebrain and to build more complete models of sexual function and dysfunction across vertebrate life.

In most fishes, amphibians, and non-avian reptiles (altogether representing a majority of vertebrate species), sex determination is driven by polygenic or even environmental factors such as temperature or pH during embryonic development (5–9). In many teleost fishes, sexual differentiation is largely accomplished through reversible actions of gonadal steroids even in adulthood (10–12). The remarkable sexual plasticity in teleost fishes has even allowed for the repeated evolution of hermaphroditism in various forms (13–16). The most common and well-studied form of hermaphroditism in teleosts is sequential hermaphroditism, or sex change. In sex-changing species, an adult fish that is actively reproductive as one sex will naturally and completely change sex when a certain age, size, or social status is acquired (13, 17). Most socially controlled sex changing species are protogynous, i.e. they initially develop as female and change sex to male if they ascend to dominance in the social hierarchy. Anemonefishes (clownfishes) are unusual in that they display the reverse pattern. All female anemonefishes were once males that changed sex to female after ascending to social dominance (18).Anemonefish are thus a model for understanding the unique case of female sexual differentiation from a male baseline in the absence of strict genetic determinism.

The remarkable behavioral and gonadal transformation during sex change in anemonefish implies a corresponding transformation of sexually-differentiated control mechanisms in the forebrain that regulate gonadal function, reproductive behaviors, and social decision-making. Indeed, many sexually dimorphic mammalian forebrain regions are thought to be conserved in teleost fishes, for example the amygdala, basal nucleus of the stria terminalis, and various hypothalamic subnuclei (19–21). Perhaps the best-known example is the sexually-differentiated POA, a key hypothalamic node of the HPG axis that regulates gonadal physiology, mating behavior and mate preferences and is well-conserved across fishes, amphibians, reptiles, birds and mammals, including humans (4, 22–26). However, research in sex-changing fishes has thus far focused on either a small number of cell types and molecules, e.g., GnRH1 neurons (27, 28), AVT neurons (29), aromatase (30–32) and isotocin (33), or has performed bulk RNA-sequencing analysis of the entire brain or multiple brain regions combined (34–36). We simply do not yet understand the neural basis of adult sex change at a cellular level. New insight into molecular mechanisms and targets of neurosexual differentiation will further enhance our understanding of this remarkable specialization in sexual biology in which the adult brain controls a complete transformation of the gonads and genitalia.

Recent advances in single cell sequencing technology are enabling researchers to describe brain sex differences with dramatically greater depth and resolution (37, 38). These tools are species-agnostic, allowing researchers to accelerate the development of research programs in non-model organisms (39–42). Here we apply single nucleus RNA-sequencing (snRNA-seq) to investigate the forebrain of the iconic sex-changing anemonefish, *A. ocellaris*. We generate a single-cell atlas of the male and female anemonefish forebrain and test for sex differences in gene expression, proportion, neuronal excitation, and predicted communication across cell types. We uncover unanticipated and extensive sex differences spanning multiple neuronal and glial populations, establishing a new framework for understanding neurobiological basis of adult sex change.

## Results

### Cellular atlas of the anemonefish forebrain

#### Clusters accurately reflect major cell classes

We placed male *A. ocellaris* anemonefish together in pairs (n=12 individuals, 6 pairs). After one member of each pair had completely transformed into a female and the pair had produced viable offspring numerous times (see Methods), we dissected the forebrain (containing the entire telencephalon and POA) for snRNA-seq. In total, we sequenced and analyzed 21,929 nuclei isolated from these 12 individuals (Fig. 1AB; see Supplementary Excel File 1 for fish body size and sequencing pool). On average, 2,026 genes and 4,322 transcripts were detected per nucleus (see Supplementary Figure 1 for complete summary). Nuclei were first clustered into 19 “parent” and, nested within those parent clusters, 48 “child” clusters based on gene expression profiles (Fig. 1B). Clusters were then assigned to one of eight major cell classes (Fig. 1C) based on expression of canonical cell type-specific marker genes (Supplementary Fig. 2).

**Figure 1.**
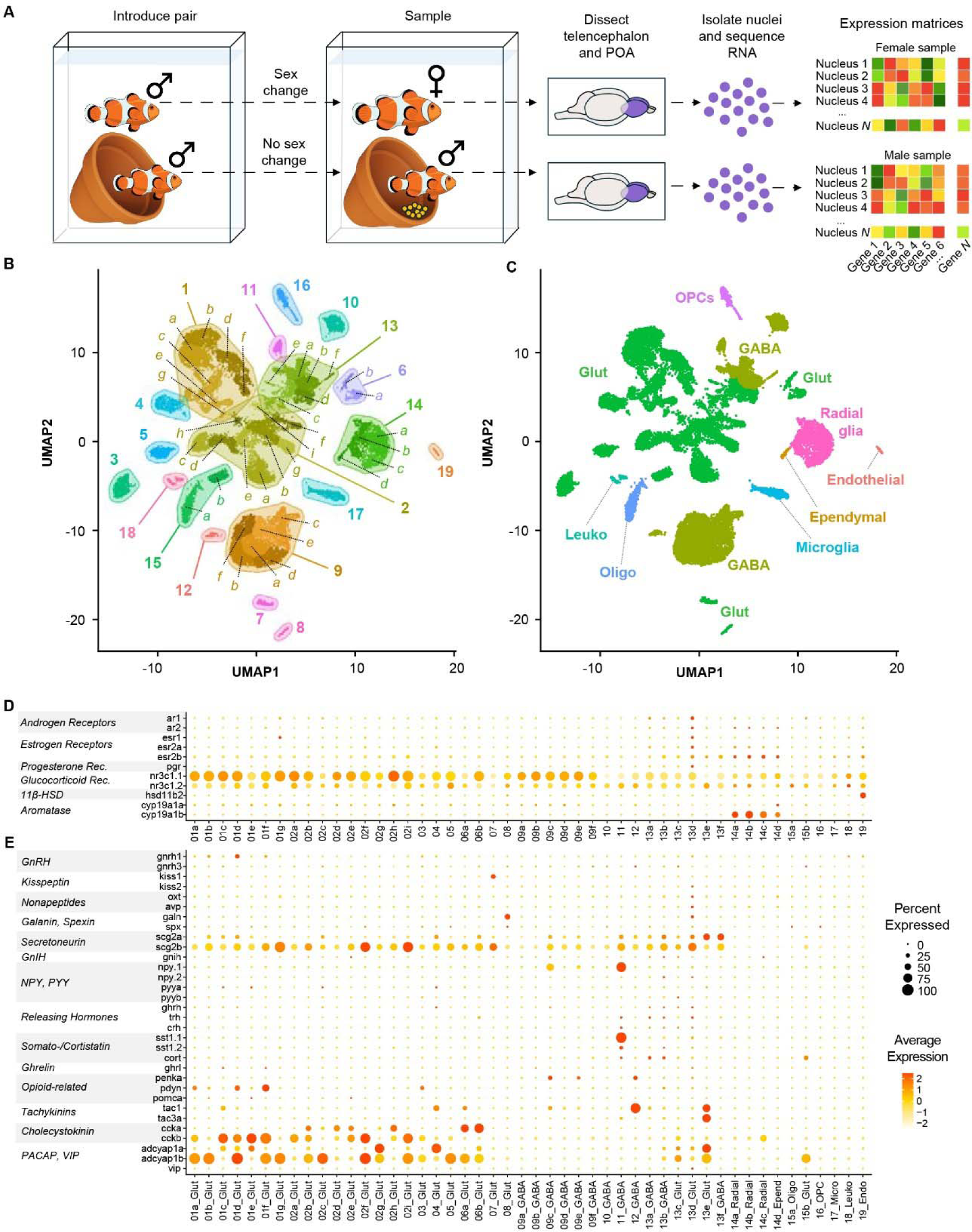
Cellular atlas of the anemonefish forebrain contains all expected cell types and identifies key cell types involved in reproduction. A) Experimental design and workflow. B) Single nuclei plotted as points in UMAP coordinate space and colored according to cluster assignment. Clusters were assigned based on transcriptomic similarity/dissimilarity with other nuclei. Parent clusters were identified with numbers (1–19), and if a parent cluster contained multiple child clusters then each child cluster was assigned a letter (a, b, c, etc.). Colored fields surround parent clusters to aid in identifying their boundaries. C) Clusters were assigned to cell types based on expression of canonical marker genes among the cluster’s marker-DEGs. For more information on marker gene expression used to assign cell types, see Supplementary Excel File 2. D-E) Dotplots depict the expression of select genes within each cluster, chosen *a priori* due to their documented involvement in the differential regulation of reproduction between sexes. Dot size indicates the percent of cells within the cluster that expressed the given gene. Dot color indicates the average normalized expression value within the cluster. These genes fall into two general categories: those involved in steroid signaling (D), and those that mark neuronal populations that release particular neuropeptides (E). Genes associated with small molecule neurotransmitters are described in Supplementary Fig 4.

Neurons were broadly classified as glutamatergic (*slc17a6a+* and/or *slc17a6b+*) or GABAergic (*slc32a1+*, *gad2+* and/or *gad1b+*), and glutamatergic and GABAergic neurons segregated strongly across clusters (Supplementary Fig. 2). Radial glia (*cyp19a1b+* and/or *gfap*+) (43, 44) were split into four child clusters, one of which (14d) strongly and selectively expressed *crocc2* (an essential cilia protein component) and was thus identified as a putative ependymal or ciliated cell population (45). Radial glial child clusters 14a-c expressed several genes involved in neurotransmitter reuptake (*slc6a11b*, *slc1a2b*, *glula*) that are expressed in mammalian astrocytes (46), supporting the hypothesis that radial glial cells in fish perform functions characteristic of mammalian astrocytes (43). Other less abundant cell classes included oligodendrocytes (cluster 15a; *mbpa+*), oligodendrocyte precursor cells (cluster 16; *cspg4+*), microglia (cluster 17; *p2ry12+*), leukocytes (non-microglial) (cluster 18; *ptprc+*/*p2ry12-*), and endothelial cells (cluster 19; *vegfd+*) (Supplementary Fig. 2). All major cell types were detected in both sexes and in all individuals in roughly similar proportions (Supplementary Fig. 3). These clusters align strongly with major known cell types in the teleost forebrain.

#### Cluster transcriptomes identify distinct neuromodulatory signaling systems

To further characterize cluster biology, we investigated expression patterns of genes associated with specific neuromodulatory systems, i.e. major small molecule neurotransmitters, neuropeptides, and steroid hormone receptors (a subset is shown in Figure 1D-E; complete results in Supplementary Excel file 2, Supplementary Figures 4-6). The analysis revealed two cholinergic (*chat*+ and/or *slc18a3b*+) neuronal clusters (7a, 13e) as well as clusters containing multiple nuclei that co-expressed the dopamine transporter and synthesis enzyme genes (*th+* and *slc6a3+*; most frequently in cluster 9f, less frequently in 13d, 13b, 13a, and 9d). Only a small number of nuclei co-expressed transporter and synthesis enzyme genes for norepinephrine (*dbh+* and *slc6a2+*; clusters 2a, 6a, and 9b) and serotonin (*tph1+* and *slc6a4b+*; cluster 13d; Supplementary Fig. 4). We speculate that the low prevalence of nuclei co-expressing both transporter and synthesis enzymes reflects the high degree of sparsity of snRNA-seq data generally, and the low recovery of *dbh, tph1*, and *slc6a4b* transcripts in particular. Several clusters were enriched for neuropeptide genes implicated in reproduction in fish, including *gnrh1*, *kiss1*, *galn*, *npy*, *oxt*, *avp*, *pdyn*, *penka*, *tac1*, *tac3a*, *ccka* and *cckb*, *scg2b*, and *adcyap1a* and *adcyap1b* (Fig. 1E; Supplementary Fig. 5). Notably, cluster 13d selectively expressed a subset of neuropeptide-related genes (*avp*, *oxt, galn*) known to be selectively expressed in the fish POA, and multiple transcription factor markers known from mouse POA (*hmx3a*, *hmx2*, *hmx3*), supporting the conclusion that nuclei in this cluster are POA-derived (47). Other POA neuropeptide genes (e.g. *crhb, trh*, *crh*) were expressed in cluster 13d as well as other clusters (e.g. clusters 1a, 1b, 11, 13a, 13b, 13c). Interestingly, *gnrh1*, a neuropeptide that is selectively expressed in the fish POA, was strongly and preferentially expressed in glutamatergic cluster 1d. One plausible explanation is that *gnrh1+* nuclei in cluster 1d are anatomically derived from the POA but are developmentally derived from a distinct cell type that does not express other POA markers *hmx3a*, *hmx2*, or *hmx3*. Ultimately, these data are consistent with recent work in zebrafish showing that the POA is populated by multiple neuronal subpopulations from distinct neurodevelopmental cell lineages (48).

In addition to monoamine and neuropeptide systems, we also investigated steroid systems associated with sex and social dominance. Sex steroids (e.g. estrogens, androgens) are key mediators of sexual differentiation across vertebrate life (4), and glucocorticoid signaling is a key mediator of dominance status between the sexes (49, 50). To identify cell populations that may respond to steroid hormone signals, we investigated cluster-specific expression patterns of sex steroid and glucocorticoid receptor genes (Fig. 1D; Supplementary Fig. 6). Putatively POA-derived cluster 13d was enriched for all sex steroid receptors (*ar1*, *ar2*, *esr2a*, *esr2b*, and *pgr*), consistent with known steroid hormone sensitivity of the POA in teleosts and other vertebrates species (4, 51, 52). Notably, all radial glial child clusters (including putative ependymal cells) were enriched for *esr2b* and *ar2*, consistent with steroid modulation of neurogenesis (53). Glucocorticoid receptor (*nr3c1*) was widely expressed and was enriched in parent clusters 1, 2, and 9 (specifically, glutamatergic 1a-d, f and g; glutamatergic 2a, b, d, e, h and i; and GABAergic 9a-f). Together these data support neuromodulatory signaling architecture as a major axis distinguishing clusters in this study and reveal a suite of candidate steroid hormone-sensitive neuronal and glial populations, including putatively POA-derived cluster 13d.

#### Gene expression supports strong anatomical delineation of clusters

Previous work in other species has shown that snRNA-seq clusters can segregate strongly along neuroanatomical dimensions (42, 54). To investigate the potential neuroanatomical origins of nuclei in our dataset, we first examined expression patterns of previously published teleost neuroanatomical marker genes (42) across clusters. This revealed a strong pattern whereby glutamatergic neuronal clusters tended to express genetic markers of the dorsal pallium and GABAergic neuronal clusters tended to express genetic markers of the subpallium, mirroring findings in other teleosts (55). Next, we employed SAMap to align anemonefish snRNA-seq clusters with previously published cichlid snRNA-seq clusters, which had been correlated to forebrain regions through integration with spatial transcriptomics. Briefly, SAMap estimates the cross-species transcriptional similarity of snRNA-seq cell types, accounting for the degree of protein sequence similarity between species (56). We observed remarkably strong correspondence between anemonefish and cichlid clusters, with most clusters being mutual best matches, a pattern that spanned radial glia, oligodendrocytes, oligodendrocyte precursor cells, microglia, and many neuronal subtypes (Supplementary Fig. 8). These results support strong conservation of forebrain cell populations across teleost species. Many glutamatergic child clusters in anemonefish were transcriptionally similar to glutamatergic clusters in cichlids that spatially mapped to specific forebrain regions (54). For example, anemonefish cluster 2 aligned strongly with cichlid cluster 10_Glut, which spatially mapped to the dorsal medial region of the telencephalon (Dm). In contrast, anemonefish cluster 3 aligned strongly with cichlid cluster 12_Glut, which spatially mapped to the granule subregion within the dorsal lateral pallium (Dl-g), a subregion that was recently shown to bear strong transcriptional similarities with the mammalian retrosplenial cortex (54). These results support a level of neuroanatomical dimensionality to the anemonefish clusters, identifying specific brain regions from which specific clusters were likely derived. This provides a plausible anatomical basis to enrich our understanding of the sex differences revealed in subsequent analyses (see below).

#### Predicted cell-cell communication between clusters identifies predominant sending and receiving populations

In the brain, heterogeneous cell populations are interconnected to molecularly communicate in functional circuits. To investigate possible patterns of molecular communication among cell populations identified here, we used NeuronChat to perform cell-cell communication analysis (57) Briefly, this analysis uses gene expression associated with known ligand-receptor interactions to estimate the molecular potential for directional communication (“connection weight”) between all possible pairs of putative “sender” and “receiver” cell populations. We analyzed predicted communication potential among parent clusters, child clusters, and gene-defined subpopulations of the putatively POA-derived cluster 13d (Supplementary Excel file 3). The connections with highest weights were dominated by GABAergic and putatively interneuronal clusters 10 (*pvalb6+*) and 11 (*sst+*) as sending populations, followed by clusters 13a, 13b, 13f, 9c, and 13d *galn+* and 13d *esr2a+* neurons. Interestingly, on average, oligodendrocyte precursor cells had the strongest receiving weights, followed by glutamatergic neuronal cluster 3. Within putatively POA-derived cluster 13d, the strongest receiver subpopulations were *ghrhrb+*, *kissr*+, and *galn+* nuclei. Overall, predicted communication was strongly driven by expression patterns of Neurexin– Neuroligin gene pairs, followed by genes associated with GABAergic signaling (Supplementary Fig. 7). These results confirm that clusters in this study are heterogeneous in predicted communication with one another. Furthermore, they provide a plausible means of inferring not just sex differences across individual clusters but across full putative neural circuits in subsequent analyses (see below).

### Sex change leads to extensive reorganization of the forebrain

#### Sex differences in cell type-specific gene expression are pervasive and pronounced

Previous work in anemonefish and other sex-changing fishes has shown sex differences in brain gene expression, but only for a small number of genes (34–36, 58). However, these studies were limited by analysis of whole brain or whole brain region homogenate rather than analysis of individual cells or nuclei. Because the forebrain contains conserved brain regions and heterogeneous cell populations that regulate gonadal physiology, reproductive behavior, and social decision-making, we hypothesized that a subset of cell populations would display robust and widespread sex differences in gene expression. In support of this hypothesis, a large suite of neuronal and glial clusters exhibited strong sex differences in gene expression (Figure 2AB; Supplementary Excel Files 4 and 5). Among parent clusters, a disproportionately large number of genes exhibited sex-associated expression (sex-DEGs) across a suite of neuronal populations (clusters 1, 2, 3, 9, 13) as well as microglia (cluster 17) (Supplementary Excel File 4). These patterns were explained in part by a set of constituent child clusters exhibiting a disproportionately large number of sex-DEGs (1a, 1b, 2a, 9a, 9c, 9d, 13a, 13b) as well as radial glial cluster 14b and mature oligodendrocyte cluster 15a (Fig. 2A,B; Supplementary Excel File 5). In gene ontology (GO) enrichment testing for biological processes, sex-DEGs were enriched for distinct functions in specific clusters (see Supplementary Results for more information). For example, sex-DEGs in cluster glutamatergic cluster 2a were uniquely enriched for “GABA-A receptor complex” and “chloride channel complex”, possibly reflecting transcriptional effects of sex differences in inhibitory input.

**Figure 2.**
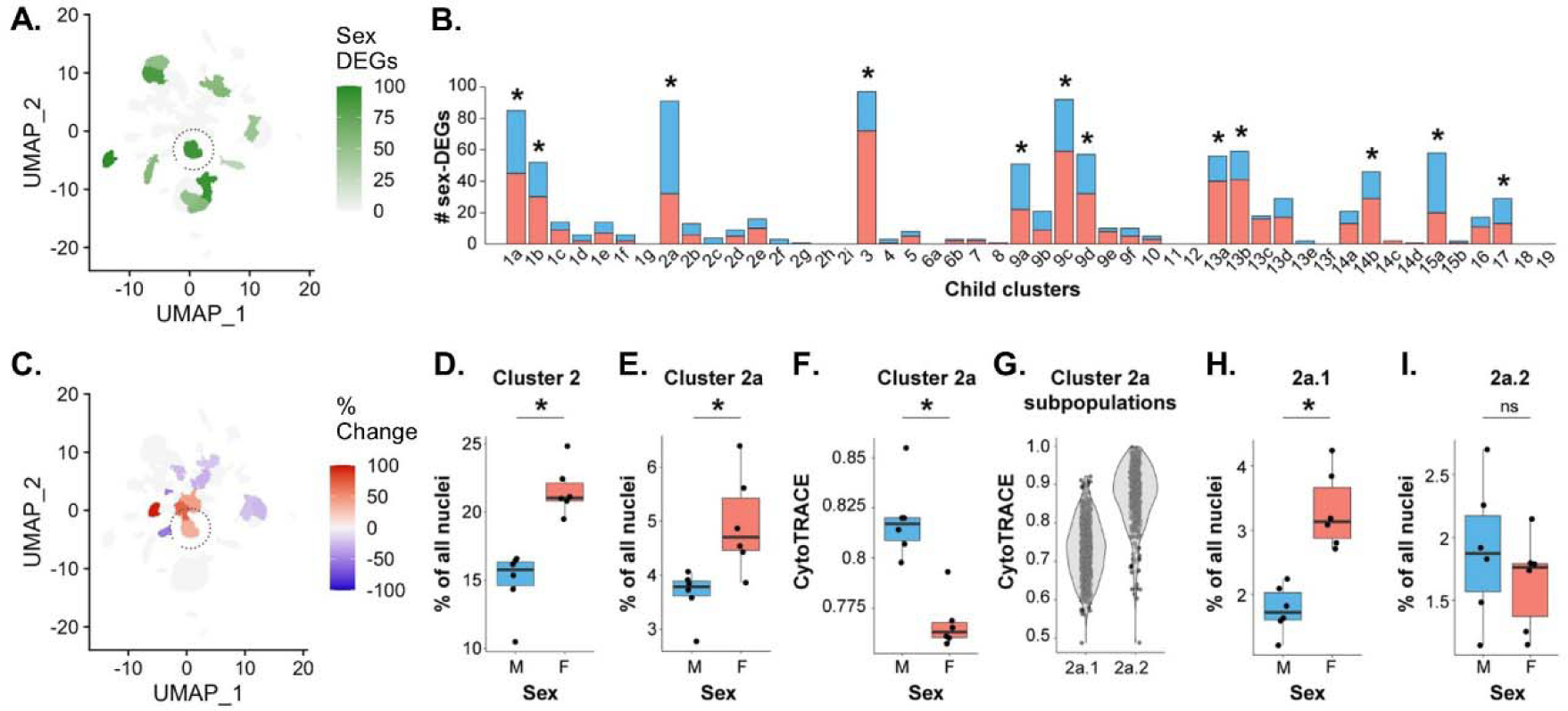
Widespread sex differences in cell type-specific transcriptomes and proportions of cells in the anemonefish forebrain. A) UMAP plot indicating the number of sex-DEGs by child cluster. Only clusters that were significantly enriched for sex-DEGs are shown. B) Bar plot showing the number of sex-DEGs per cluster. Asterisks indicate clusters that were significantly enriched for sex-DEGs, also shown in panel A. Red filled portions of the bars show the number of sex-DEGs that were upregulated in females compared to males. Blue filled portions show the number of sex-DEGs that were upregulated in males compared to females. C) UMAP plot showing sex differences in cluster proportions. Sex differences in proportion are presented as percent change from male to female. Only clusters that showed a significant proportion difference (q < 0.05) are colored. Eleven of 48 clusters (representing 27% of all cells) differed in proportion between sexes. D-E) The percentage of nuclei in parent cluster 2 and child cluster 2a in individual males and females are shown, females displayed higher percentages than males. F) In cluster 2a, average CytoTRACE scores (across all nuclei within individuals) were greater (indicating relatively less mature) in males than females. G) Within 2a, CytoTRACE scores indicated 2a.1 as the more differentiated/mature subcluster and 2a.2 as the less differentiated and more immature subcluster within 2a. H) The relatively mature subcluster 2a.1 was approximately twice as abundant in females than males. I) The relatively immature subcluster 2a.2 was similar in abundance between males and females. * above the bars indicates significance at q<0.05.

Several clusters showed strong directional biases in the proportions of sex-DEGs that were downregulated versus upregulated in females versus males (Fig 2B; Supplementary Excel Files 4 and 5). More sex-DEGs were upregulated in females in glutamatergic neuronal parent clusters 3 and 13 (and a similar trend was observed in radial glial cluster 14); and more sex-DEGs were downregulated in females in glutamatergic neuronal parent clusters 1, 2, and 15. These patterns were mirrored in a set of constituent child clusters (2a, 13a, 13b, 13c), and were also observed in GABAergic cluster 9c, which had more sex-DEGs upregulated in females versus males. Enrichment testing revealed that sex-DEGs that were upregulated in females versus males (and vice versa) were related to specific biological functions (Supplementary Excel Files 4 and 5). For example, in parent clusters 1, 2, and 15, sex-DEGs that were upregulated in females were differentially enriched for GO categories related to apoptosis (clusters 2 and 15) or autophagy (cluster 1), whereas sex-DEGs that were downregulated in females were differentially enriched for multiple categories related to neuronal differentiation (Supplementary Results).

#### Sex differences in cellular composition of the forebrain are widespread and prominent

Previous work in teleosts (including anemonefish) and mammals identify sex differences in the relative proportions of specific cell populations in the forebrain, including within the POA (4, 27, 28, 38, 59, 60). Based on this work, we tested for sex differences in the relative proportions of clusters and putatively POA-derived cell populations. A number of both neuronal and glial populations showed sex differences in relative proportions (Fig. 2C, complete results for parent clusters, child clusters, and POA-derived populations shown in Supplementary Excel Files 8-10, respectively). Among parent clusters, sex differences were observed in glutamatergic neuronal cluster 2 (more abundant in females) and radial glial cluster 14 (more abundant in males) (Supplementary Excel File 8). Among child clusters, the sexes differed in the relative proportions of multiple glutamatergic neuronal clusters (1f, 1g, 2a, 2b, 2c, 2e, 13c and 15b), GABAergic cluster 13a, and radial glial clusters 14a and 14b (Fig. 2C; Supplementary Excel File 9). Taken together, these results support the idea that sex change leads to widespread differences in the neuronal and glial composition of the anemonefish forebrain, with 11 of 48 child clusters (comprising about 27% of all cells) being sexually differentiated in cell abundance.

#### Cluster 2 stands out as strongly differentiated in gene expression and cell abundance

Glutamatergic neuronal parent cluster 2 (Fig. 2D), and several embedded child clusters (2a, b, c and e) were 35-95% more abundant in females versus males (Supplementary Excel Files 8 and 9). Cluster 2 also had a disproportionately large number of sex-DEGs, a disproportionately large number of female-downregulated sex-DEGs, and sex-DEGs were enriched for biological processes related to neuronal differentiation and apoptosis (Fig. 2B, see “Cell type-specific gene expression” above). Among its constituent child clusters, sex differences in proportions were the most pronounced in 2a (Fig. 2E), which preferentially expressed *cckb*, the gene encoding cholecystokinin, and *galr2a*, the gene encoding galanin receptor 2. To explore cluster 2a in more detail, we investigated sex differences in neuronal maturity using CytoTRACE, a bioinformatics tool that estimates cellular maturity based on the gene expression. Cluster 2a nuclei were predicted to be relatively more mature in females compared to males (Fig. 2F; cluster 2a was the only child cluster with a sex difference in CytoTRACE score; Supplementary Excel File 11; 0 indicates high relative maturity, 1 indicates low relative maturity). To better understand this pattern, we re-clustered 2a independently from other nuclei and identified two subpopulations of 2a that differed in relative maturity: cluster 2a.1 was predicted to be relatively more mature and 2a.2 was predicted to be relatively less mature (Fig. 2G). Females displayed approximately double the proportion of the relatively mature cells in subcluster 2a.1 compared to males (Fig. 2H), but similar proportions of the relatively immature cells in subcluster 2a.2 (Fig. 2I). The relatively mature subpopulation expressed both *cckb* and *gal2ra*. These data support a model in which protandrous sex change involves maturation and increased abundance of a population of glutamatergic, galanin-sensitive, CCK-secreting neurons in the Dm, while maintaining consistent proportions of the relatively immature subpopulation. CCK signaling is known to play a role in the neural control of reproduction in fish and a recent study in zebrafish shows that CCK signaling is required for the development of female gonads (51, 61).

#### Sex change involves substantial reorganization of the steroid-sensitive and neuropeptidergic subpopulations of the POA

Given their established roles in regulating sexually-differentiated physiology and behavior in anemonefish (62–64) we hypothesized that steroid-sensitive and neuropeptide-synthesizing neuronal populations would also exhibit sexually dimorphic proportions and/or transcriptome. To investigate this, we tested for differences in the proportions of nuclei expressing specific steroid hormone receptors, neuropeptides or neuropeptide receptors known to be involved in the neural control of reproduction, specifically within putatively POA-derived cluster 13d (Supplementary Excel File 10). In support of our hypothesis, we found males displayed a significantly greater proportion of nuclei expressing *adcyap1a, esr2a* (Fig. 3A)*, npy7r,* and *tacr3,* whereas females displayed a greater proportion of nuclei expressing *esr2b* (Fig. 3B), *cckbr2* (Fig. 3C), and *pgr* (Fig. 3D). Notably, *esr2b* and *pgr* were strongly co-expressed across nuclei (see Supplementary Figure 9), suggesting that increased proportions of *pgr*+ and *esr2b*+ in females may be driven by the same underlying cell population either increasing in abundance or, alternatively, upregulating *esr2b* and *pgr*. Taken together, these results support the idea that transformation from male to female is associated with substantial molecular and cellular reorganization within the preoptic area, including differentiation of signaling systems known to regulate reproductive physiology in other teleosts and plausibly playing a similar role in anemonefish (e.g. estrogen, cholecystokin). Such changes may support the transition from male to female and/or maintenance of sex-specific physiology and behavior.

**Figure 3.**
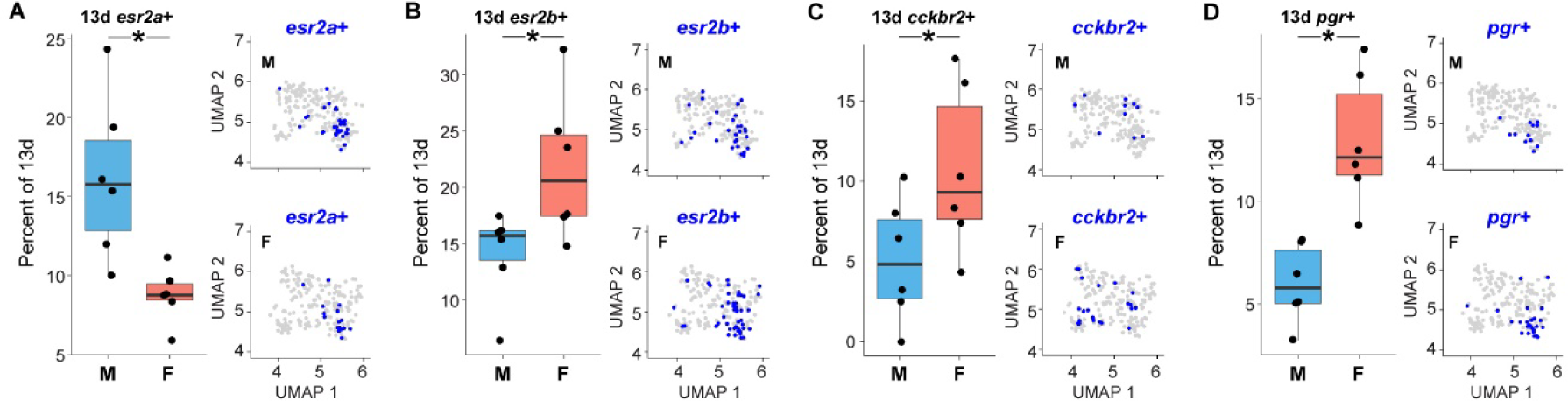
The putative POA-derived cluster contains sexually-differentiated subpopulations marked by receptors for estrogen, progesterone, and CCK. Boxplots show sex differences in proportions of sex steroid receptor expressing neurons in cluster 13d that map to the POA. A) Males display a greater proportion of cells in 13d that are *esr2a*+ than females. UMAP plot of cells in cluster 13d, cells colored blue if they express *esr2a*. B-D) Females display a greater proportion of cells in 13d that are *esr2b+*, *cckbr2+,* and *pgr+* as compared to males. * above the bars indicates significance p<0.05.

#### Sex differences in radial glia are correlated with specific neuronal sex differences

In adult teleosts, radial glial cells supply new neurons throughout the brain and are critically involved in the differentiation and maturation of progenitor neurons (11, 42–44, 53, 65). In our earlier analyses, radial glial clusters 14a and 14b were more abundant in males, and cluster 14b contained a disproportionately large number of sex-DEGs (see above). Earlier analyses also identified sex differences in neuronal proportions and expression of genes related to neurogenesis. Given these findings together, we hypothesized that radial glia play a role in sex change and/or maintenance of sex-specific phenotypes. To test this hypothesis, we re-clustered radial glia independently of all other nuclei and identified nine subclusters. These subclusters differed in transcriptional signatures of cell maturity, radial glial functional states, and expression of notable marker genes including aromatase (strongly expressed in subclusters 2, 4, 5 and 6; Supplementary Figure 10, Supplementary Excel Files 12 and 13). Further analysis suggested sex differences in multiple dimensions of radial glial biology. Radial glial subclusters 0 and 3 displayed a disproportionately large number of sex-DEGs (Supplementary Excel File 14; Supplementary Figure 11). The composition of specific subclusters also differed between the sexes, with females exhibiting greater proportions of the relatively immature subcluster 8 and males exhibiting greater proportions of the relatively mature subcluster 5 (Supplementary Excel File 15). Females showed stronger expression of aromatase compared to males in subcluster 4 (Supplementary Excel File 14), mirroring whole-brain patterns in *A. ocellaris* and other species (34, 58, 66, 67).

We next hypothesized that radial glia may play a role in producing some of the neuronal sex differences identified above. To test one aspect of this, we asked whether radial glial proportions or aromatase expression were correlated with proportions of particular neuronal populations. Indeed, the relative proportions of radial glial subclusters 0 and 5 were strongly and inversely correlated with the proportions of a subset of neuronal populations that differed in abundance between the sexes (relatively mature 2a, 2c, 2e, 13c, 15b, 1g), and multiple putatively POA-derived subpopulations (13d *tacr3*+, 13d *adcyap1+,* 13d *pgr+*; see Supplementary Figure 12). Second, aromatase expression in radial glial subcluster 4 was positively correlated with the relative proportions of 13d *esr2a+,* 13d *esr2b+*, 13d *npy7r+* and 13d *cckbr2+* nuclei. Taken together, these results are consistent with aromatase expression in a specific subpopulation of radial glia being related to reorganization of estrogen and CCK signaling systems in the POA, and more broadly in which distinct radial glial subpopulations are positioned to drive distinct components of sexual differentiation throughout the forebrain.

#### Sex change leads to substantial differences in excitability of multiple neuronal populations

Differential regulation of the gonads and behavior in males versus females could arise from differences in the excitability/activation of specific neural circuits. To test for possible sex differences in neuronal activity we analyzed expression of immediate early genes (IEGs) across clusters and putatively preoptic neuronal subpopulations. Fish in this study did not receive any specific stimulus or display any particular behavior immediately prior to tissue collection, therefore any consistent sex differences in IEG expression may best reflect sex differences in baseline activity. Because individual IEGs are recovered at relatively low levels in snRNA-seq data, we followed the same approach as Johnson et al. (42) and tracked a set of 22 genes, each of which was selectively co-expressed with *egr1*, *npas4a*, and *fosab* independently across many neuronal clusters (see Methods). Sex-associated IEG expression was only observed in neuronal clusters, including four parent clusters (3, 9, 12, and 13; Supplementary Excel File 16) and five child clusters (2b, 9c, 13a, 13b, and 13d; Fig. 4; Supplementary Excel File 17). Within putatively POA-derived cluster 13d, seven subpopulations showed sex-associated IEG expression (13d *oxt*+, *npy*+, *pdyn*+, *scg2b*+, *tac1*+, *vipr1a*+; Supplementary Excel File 18). Notably, all of these populations had strong predicted potentials to receive communication from clusters 3, 12, 9c, 13a, and 13b (Supplementary Excel File 3), which were all the specific clusters identified above as having significant sex differences in IEG expression (Supplementary Excel File 17). Strikingly, IEG expression was greater in females versus males in every case except for child cluster 2b (Fig. 4A). This overarching pattern may reflect broadly increased excitability across a suite of neuronal populations in females compared to males. One possible explanation is that males and females have a different circulating steroid milieu, which can regulate neuronal excitability in a cell-type dependent manner (68–71). Nonetheless, these results support a model in which adult sex change leads to increased excitability of multiple neuronal populations in the forebrain that are synaptically interconnected and communicating with one another, including steroid hormone-sensitive neuronal populations in the POA that regulate reproductive physiology and behavior. Together with earlier analyses, these results highlight a suite of neuronal populations (clusters 2a, 2b, 3, 9c, 13a, 13b) as sexually differentiated across multiple lines of analysis.

**Figure 4.**
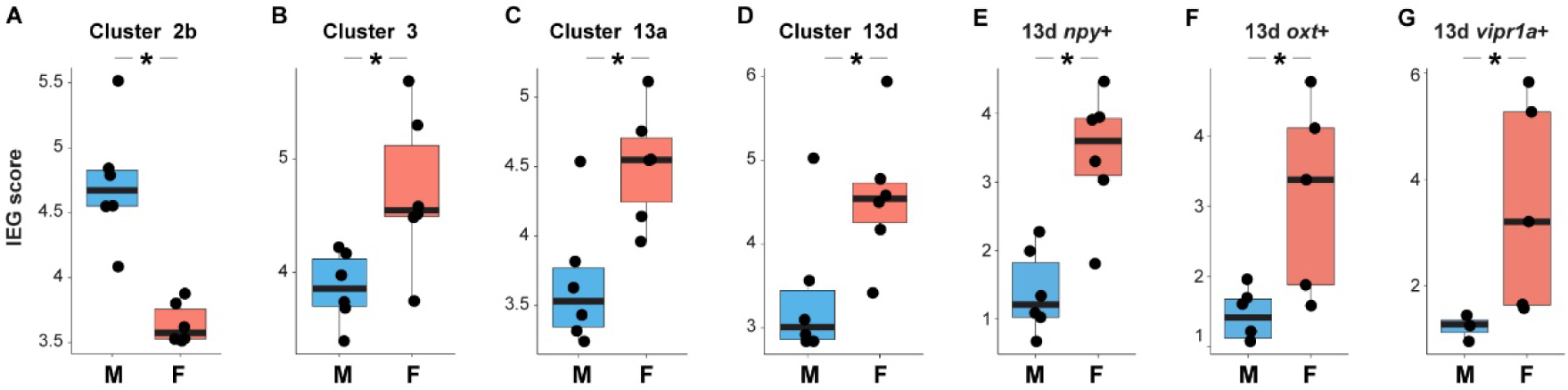
Sex differences in IEG expression across clusters including putatively POA-derived subpopulations; most sexually differentiated clusters showed higher IEG expression in females. A) Males display a higher IEG score (see methods-Sex Differences in IEG Expression) than females in glutamate neuron cluster 2b. B-G) Females display a higher IEG score than males in glutamatergic neuronal cluster 3, GABAergic neuronal cluster 13a, glutamatergic neuron cluster 13d, and within 13d, *npy*+, *oxt*+, and *vipr1a*+ neurons. Asterisks indicate significance (q<0.05).

#### Sex change leads to sexually dimorphic patterns of molecular communication

Results above indicated sex differences in activity levels across a specific set of clusters, many of which were also found to be highly interconnected in the initial global NeuronChat analysis (see above). To further investigate sex differences in communication among neuronal populations, we first estimated predicted communication between neuronal populations within each individual, and then used permutation testing to identify observed sex differences that were greater than 95% of shuffled differences. A disproportionately large subset of neuronal population pairs exhibited sex differences in predicted communication (p_perm_<0.05, n=49, ∼7.3%; Exact binomial test, p=0.010). Communication between sexually dimorphic clusters 2a, 2b, 3, 9c, 13a, 13b accounted for a disproportionate number of these differences (6/49 hits, Fisher’s Exact test, p=0.0081, Odds Ratio=4.21, 95% C.I.=[1.31,11.63]). For example, in females compared to males, the subpopulation within cluster 2a that was relatively more mature and more abundant in females was predicted to receive stronger communication from sexually dimorphic cluster 13a (Fig. 5). Additionally, among pairs exhibiting sex differences in predicted communication, specific neuronal populations were overrepresented as senders (13d *ar+*, 13d *esr2a+*) and receivers (clusters 2f, 3, 6b; 13d *cckbr1+*). For example, the putatively POA-derived cluster 13d *cckbr1+* subpopulation was predicted to receive stronger communication from many sender populations in females compared to males, including sexually dimorphic cluster 13a, sex-DEG enriched GABAergic neuronal clusters (9a, 9d), and clusters 9e, 9f and 10. Taken together, these results support the idea that adult sex change leads to sexually dimorphic molecular communication patterns among a specific subset of neuronal populations in the forebrain, particularly between sexually dimorphic cell populations and those expressing sex steroid receptors such as *ar*, *esr2a*, and *cckbr1*.

**Figure 5.**
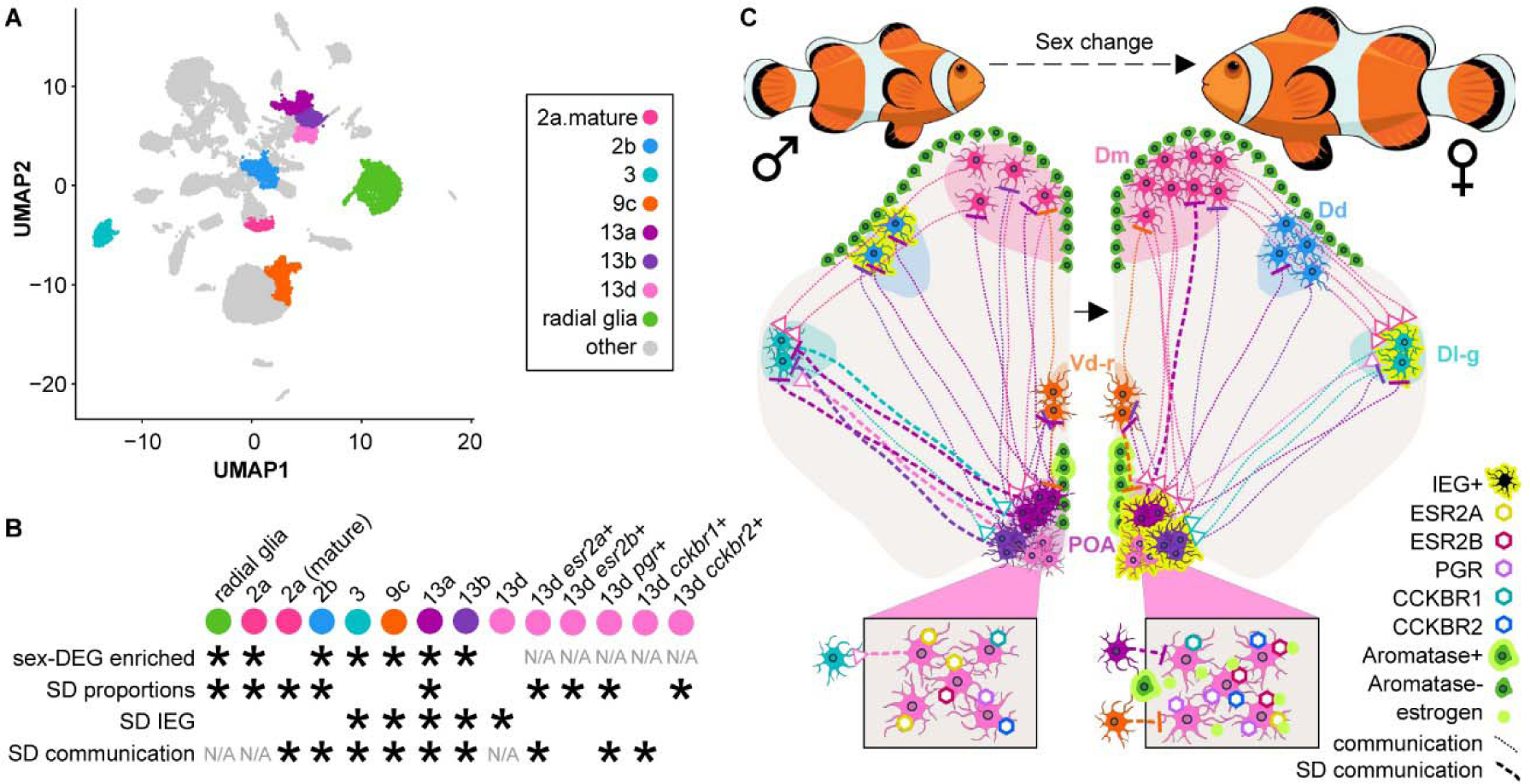
A proposed cellular axis of neurosexual differentiation. (A) UMAP plot highlighting high-confidence sexually dimorphic populations of interest that were identified by multiple lines of analysis. Color-coding scheme applies to all panels (including the color of illustrated cells in C). (B) Summary of cell type-specific sex differences in gene expression (top row), relative proportions (second row), IEG expression (third row), and predicted intercellular communication (bottom row). SD stands for sexually differentiated. Asterisks indicate populations that were identified as significant for each analysis, and N/A indicates populations that were not analyzed in particular analyses. (C) A hypothesized model of neurosexual differentiation during adult sex change in the anemonefish presented on a drawing of a coronal section of the telencephalon. The left hemisphere in this image represents the state of the male forebrain, and the right hemisphere represents the state of the female. Cell populations are drawn in hypothesized neuroanatomical locations based in part on cross-species analyses with a previously published cichlid snRNA-seq and spatial transcriptomics telencephalic atlas.

#### Sex change leads to a sexually differentiated neurobiological axis

Analyses of sex differences in gene expression, cell type-specific proportions, signatures of neuronal excitation, and cell-cell communication converged on a set of major neuronal hubs for sex differences in the anemonefish forebrain (clusters 2a-b, 3, 9c,13a-b, and the putatively POA-derived cluster 13d+ *esr2a+* subpopulation). Cross-species mapping to cichlid snRNA-seq clusters supported a subset of specific forebrain regions from which these clusters were likely derived, including Dm (2a), Dd (2b), Dl-g (3), Vd-r (9c), and POA (13a, 13b, 13d *esr2a*+). Tracing studies in other fish species have demonstrated strong connectivity among these regions (72–74), consistent with the strong predicted cell-cell communication patterns among these sexually dimorphic neuronal hubs. Additionally, overrepresentation of these high-confidence sexually dimorphic neuronal populations among predicted sex differences in communication (particularly between 2a, 3, 13a, and 13b) further supports the idea that together these populations represent a cohesive, integrated, sexually-dimorphic neural network spanning multiple forebrain regions, including the POA. In addition to these neuronal populations, multiple lines of analysis supported sexual dimorphisms in radial glia subpopulations and the estrogen and CCK signaling systems. Thus, we propose a model in which interaction among multiple subpopulations of radial glia and neurons in the forebrain, as well as estrogen and CCK synthesis and signaling among these populations, together represent a high-confidence and functionally integrated axis of neurosexual differentiation during adult sex change (Fig. 5).

## Discussion

Here we provide the first cellular atlas of the forebrain in a sex-changing fish. We uncover extensive divergence between males and females in cell type-specific gene expression, relative proportions, signatures of neuronal excitation, and predicted intercellular communication. Converging lines of evidence supported a central neurobiological axis for sex differences including radial glia, estrogen and cholecystokinin signaling systems, and a suite of interacting neuronal subpopulations embedded in the POA as well as multiple unexpected forebrain regions (Dm, Dd, Dl-g, and Vd-r). Taken together, our results suggest that adult sex change leads to extensive molecular and cellular reorganization of the forebrain. This work defines a set of molecular and cellular endpoints that can be traced over the course of sex change.

This study builds on a large body of work documenting sex differences in the brains of other vertebrates (75). Among fish, sex differences have been documented in brain concentrations of hormones, expression of multiple neuropeptide and steroid synthesis genes (e.g. aromatase), and levels of neurogenesis in specific brain regions (60). In sex changing fishes specifically, previous work has focused largely on protogynous and bi-directional sex changing species (28–30, 32, 33, 35, 76, 77). Far fewer studies have investigated brain sex differences in protandrous species. To our knowledge, only one other study has compared brain transcriptomes between sexes in anemonefish. That study used whole-brain bulk RNA-sequencing in a different species, *A. bicinctus* (34), and found that fish in the middle of sex change had many DEGs when compared to males or females, but only five sex-DEGs distinguished stable males from stable females (34). Another recent bulk RNA-sequencing study in a bi-directional sex changing goby similarly found only nine sex-DEGs across the whole brain (36). This contrasts with the extensive sex differences we found using snRNA seq. It is possible that many sex differences are cell type-specific and are washed out when cell types are pooled and analyzed together. The widespread sex differences we found across many genes and cell populations, underscores single-cell functional profiling as a powerful tool for uncovering cell type-specific sex differences and novel cell populations of interest.

The CCK system emerged as a central sexually dimorphic molecular pathway in the anemonefish forebrain. Interestingly, the forebrain CCK system is regulated by gonadal hormones and modulates opposite-sex odor processing in rodents (78–80), and CCK signaling is critical for pituitary FSH activity and female sexual development in zebrafish (61). In the present study, most sexually dimorphic clusters strongly and preferentially expressed the CCK ligand gene *cckb* (relatively mature 2a, 2b, 3) or CCK receptor genes *cckbr1* (9c, 13a) and *cckbr2* (3, 9c). Furthermore, the putatively POA-derived cluster 13d *cckbr2*+ subpopulation was significantly more abundant in females and co-expressed *cckb.* The *cckbr1*+ subpopulation was overrepresented among sex differences in predicted cell-cell communication, with female populations predicted to be stronger receivers than male populations in every case (including from GABAergic sexually dimorphic cluster 13a). It is intriguing to speculate that CCK+ neurons in Dm (2a) and/or Dl-g (3) regulate CCKR+ neurons within the POA that in turn regulate the pituitary gonadotrophs (51, 61). Plausible pathways for such regulation were supported by cell-cell communication analysis: the strongest predicted neuronal target for glutamatergic sexually dimorphic cluster 2a was glutamatergic sexually dimorphic cluster 3, followed by putatively POA-derived glutamatergic 13d *pdyn+* and sexually dimorphic GABAergic cluster 13a. This is consistent with demonstrated projections from dorsal regions of the telencephalon to the POA in other teleosts (72–74, 81). In turn, the strongest predicted neuronal targets for glutamatergic sexually dimorphic cluster 3 included the putatively POA-derived glutamate cluster 13d *ghrhrb+* subpopulation (top predicted target) and putatively POA-derived sexually dimorphic GABAergic cluster 13a. These data raise the possibility that anemonefish recruit conserved sexually dimorphic CCK pathways during sex change, and define a set of testable interactions by which proliferation and excitation of CCK+ neurons in the dorsal pallium may regulate the POA and downstream pituitary FSH cells to promote female gonadal development during the remarkable transition from male to female.

Galanin and progesterone also emerged as candidate signaling systems linking the POA to sexually dimorphic glutamatergic cluster 2a and GABAergic cluster 9c. Predicted cell-cell communication from the putatively POA-derived cluster 13d *galn*+ subpopulation to the relatively mature 2a subpopulation was among the strongest of all communications analyzed by NeuronChat, and the relatively mature 2a subpopulation nearly exclusively expressed the galanin receptor gene *galr2a*. Recent work has demonstrated galanin’s influence on reproductive behavior and physiology in fish, including alternative mating strategies of male fish and paternal care behavior (82–89). Thus, sex differences in galanin signaling could contribute to the strong sexual dimorphism in parental care observed in anemonefish, with males doing the vast majority of egg care (90). In females compared to males, putatively POA-derived cluster 13d *pgr*+ neurons were more than twice as numerous (Fig. 3D, no overlap between sexes) and were predicted to receive stronger communication from sexually dimorphic and GABAergic cluster 9c. Previous work has shown that progesterone receptor signaling is critical for sexual differentiation of the POA in mammals, but little is known about its role in sexual differentiation and function of the POA in fish. Similar to mammals, progesterone in the brain of zebrafish has two sources: gonad-derived in systemic circulation, and locally-synthesized in the brain (91). Further, progesterone receptor *pgr* is upregulated by estrogens in the brain of adult zebrafish (92) as it is in mammals (93, 94). Thus, higher brain aromatase and estradiol levels in female *A. ocellaris* (90) could account for elevated *pgr* expression relative to males. Future studies are needed to understand the functional significance of steroid and neuropeptide system reorganization within the POA during sex change, and how it relates to female reproductive biology.

Consistent with expectations, converging evidence in our results here supported strong sexual dimorphism of the estrogen system in the anemonefish forebrain. Females displayed greater expression of brain aromatase, the enzyme that synthesizes estrogen, in a specific subpopulation of radial glia compared to males, consistent with higher levels of brain estrogen in females and mirroring previous work in other species. Sex-DEGs in multiple clusters, including sexually dimorphic cluster 2a, were enriched for putative estrogen response elements, supporting the idea that estrogen contributes to sex differences cell type-specific gene expression. Perhaps most striking was the opposite proportion differences in putatively POA-derived cluster 13d *esr2a*+ and *esr2b*+ subpopulations, where males displayed a greater proportion of *esr2a*+ while females displayed a greater proportion of *esr2b*+ nuclei. These differences, as well as increased proportions of putatively POA-derived *cckbr2+* nuclei in females, were strongly correlated with one another and with aromatase expression in the radial glial subpopulation mentioned above. These results are consistent with previous studies in medaka fish, in which females also display increased expression of *esr2b* in the telencephalon and preoptic area compared to males. In medaka fish, increased *esr2b* is regulated by sex steroid hormones and is required for female mating behavior and sexual preference (95). Additional evidence in medaka suggests *esr2a* expression in the POA plays a key role in E2 feedback regulation of pituitary FSH secretion and oviduct formation (96). We speculate that a similar organization is present in anemonefish, and the rebalancing of *esr2a*+ and *esr2b*+ expression in the POA reported here is likely regulated by sex hormones, and is critical for sex-specific gonadal development and reproductive behavior.

Of the sex differences documented here, the changes in the proportions of specific cell populations are the most striking as they suggest sex change may not only involve changes in gene expression, but also a more widespread cellular reorganization of the forebrain. Consistent with these results, we have previously documented greater neurogenesis and more neurons in the anterior POA during sex change (97). Hence, neurogenesis likely contributes to increased numbers of cells in the POA cluster observed in females (Fig. 3, *esr2b*, *pgr*, *ckkbr*). Likewise, males had greater proportions of radial glia, which facilitate neurogenesis and terminal differentiation of neurons throughout adulthood in fish (98–101). This abundance of radial glia may function as a progenitor cell pool, waiting to supply those neuronal populations that are greater in females. Neurogenesis during sex change may also contribute to the large increase in the proportion of relatively mature glutamate neurons observed in cluster 2a in females (Fig. 2E). Within this cluster, males had similar numbers of relatively immature neurons as females but far fewer relatively mature neurons (Fig. 2F-I). We speculate that during sex change, some of the immature neurons in males terminally differentiate into mature neurons, while the immature population is replenished via neurogenesis to accommodate continued growth. Interestingly, recent snRNA-seq experiments suggest that neurons in the human amygdala can remain in an immature phase of development for years, and only continue maturation in response to new cues or developmental stages like puberty (102, 103). This may be analogous to the nature of sex change in anemonefish, in which the transition from male and commitment to female is the terminal and irreversible final phase of their life history. In this framework, a specific cell type in the male anemonefish brain may remain in a state of arrested development, awaiting cues to initiate terminal differentiation.

Cross-species mapping supported the conclusion that the high-confidence molecular and cellular sexual dimorphisms described above are anatomically positioned across multiple nodes of the conserved vertebrate social decision-making network (19). This network includes hormone-sensitive forebrain regions that regulate reproductive and social behaviors across species. It is thought that evolutionary and functional plasticity within this network are major mechanistic drivers of social behavioral diversity in nature (104). The social decision-making network directly interfaces with the HPG axis through the POA. Based on these results, we hypothesize that functional specializations at this interface enable reproductive adult anemonefish to change sex in response to social stimuli. Within this framework, we propose 1) reorganization of the neuromodulatory POA cell populations mediated in part by subpallial radial glial aromatase and estrogen, 2) expansion of a CCK+ glutamatergic neuronal population in Dm, and 3) altered interaction among neuronal populations in the POA, Dm, Dl-g, and Vd as high-confidence sexual dimorphisms positioned around this interface (Fig. 5). These and other predictions supported by our data will stimulate progress in our understanding of the remarkable biological innovation of adult sex change.

It is intriguing to speculate that one or more of these dimorphisms, in turn, is intertwined with the neural circuitry of social dominance. Expression of brain aromatase in subpallial radial glia is one plausible candidate. In male cichlids, social dominance is associated with decreased aromatase expression in the preoptic area (105). In contrast, brain aromatase expression in specific forebrain radial glial subpopulations increases in association with sex-specific courtship behavior in male cichlids (the building of nest-like bower structures) and regulates gonadal mass, (42). In the sex-changing anemonefish *Amphiprion bicinctus*, dominant and actively sex-changing individuals upregulate brain aromatase before upregulating gonadal aromatase and long before sex change is complete (34). Extending this idea to *Amphiprion ocellaris* and based on our results, a functional specialization in radial glia in the preoptic area may facilitate upregulation of brain aromatase in response to the perception of dominance, initiating a feminization cascade in the brain, behavior, and gonadal physiology.

## Conclusion

Adult sex change is one of the most remarkable examples of physiological transformation and neural plasticity in adult vertebrates. Cellular profiling of the sex-changing anemonefish forebrain revealed extensive differences between males and females spanning specific molecular signaling systems, neuronal and glial subpopulations, and brain regions. Resolving the full sequence of neuromolecular changes, and determining which of these changes regulate the physiological components of sex change versus the maintenance of downstream female-specific phenotypes, are primary targets for future study. Our results establish an important foundation for future work by defining a set of molecular and cellular beginning and endpoints that can be tracked through the full course of sex change. Defining and interrupting this sequence will reveal the neural drivers of specific dimensions of sex change, improving our understanding of neural regulation and functional plasticity of the HPG axis across vertebrate life.

## Materials and Methods

### Animals and Husbandry

Anemonefishes (genus *Amphiprion*) are a highly specialized group of marine fishes with many notable characteristics including sex change, a strict social dominance hierarchy, high levels of male parental care, and symbiosis with their host anemone (41). Our species, *Amphiprion ocellaris*, is phylogenetically basal among anemonefishes (106). Sex change in anemonefish always proceeds from male to female and is under social control (18, 59, 97, 107, 108). In anemonefish groups, the breeding female is the dominant individual, and her male breeding partner is the next most dominant. When the dominant female is lost from a group, the reproductive male naturally ascends to the dominant position in the hierarchy, and this transition in social status triggers sex change. The literature describing various aspects of sex change in anemonefish is rich and extends back decades (27, 59, 97, 107–115).

Fish were bred in-house from fish originally obtained from Ocean Reefs and Aquariums (Fort Pierce, FL). Six breeding pairs of *A. ocellaris* were made by pairing juvenile males together. The larger of the two males changed sex within 6 months to a year of pairing. The reproductive pairs were left alone to spawn numerous times over several years before they were sampled for this study. All 6 pairs used in this study were observed spawning with fertilized eggs within one month prior to tissue collection. Fish were housed in twenty-gallon tall (24” x 12” x 16”) aquarium tanks integrated with a central circulating filtration system. Conditions mimicked the *A. ocellaris* natural environment (system water temperature range 26-28 C, pH range of 8.0-8.4, specific gravity of 1.026, 2:12 photoperiod with lights on at 0700 and off at 1900 h). Each tank contained one terra-cotta pot (diameter 6”) as a nest site and spawning substrate. Fish were fed twice daily with Reef Nutrition TDO Chroma Boost pellets. Experimental procedures were approved by the University of Illinois Institutional Animal Care and Use Committee.

### Tissue Collection

Fish were captured from their tanks and euthanized by rapid decapitation. All tools, surfaces, and gloves were clean and treated with RNase Zap (ThermoFisher) prior to use. The brain was removed and placed in chilled dissection medium, consisting of Hibernate AB complete medium (with 2% B27 and 0.5 mM Glutamax; BrainBits, LLC) with 200 U/mL of Protector RNase Inhibitor (Roche). Remaining submerged in the dissection medium, the brain was then microdissected with reference to a high-quality brain atlas for the cichlid fish *Astatotilapia burtoni* (Maruska Lab, Louisiana State University). The cerebellum and optic tecta were first removed, then everything caudal to the posterior commissure was removed, leaving the diencephalon portion containing preoptic area (POA) intact as well as the complete telencephalon. This dissection method was designed to prioritize speed and consistency while still preserving the POA in full. The tissue was placed in a clean RNase-free tube and stored at -80° C until nuclei isolation.

### Nuclei Isolation

For nuclei isolation and sequencing we created two pooled samples for each sex, each pool containing tissue from three individuals. Individual-level nuclei assignments were later recovered using a computational approach (see below). Samples were pooled in 5 mL LoBind tubes (Eppendorf) in 1 mL chilled lysis buffer (10 mM Tris-HCl, 10 mM NaCl, 3 mM MgCl2, and 0.1% Nonidet P40 Substitute in nuclease-free water) and incubated on ice on a shaker for 30 min. Next, 1 mL Hibernate AB complete medium was added to the tube and the tissue was triturated by repeated aspiration through a sterile silanized glass pipette (BrainBits, LLC) until the solution was homogenous. Samples were then transferred to clean 2 mL LoBind tubes for centrifugation for 5 min at 600 rcf at 4 °C. Supernatant was carefully discarded without disturbing the nuclei pellet, then 600 uL of wash buffer (2% BSA and 0.2 U/uL RNase Inhibitor in 1X PBS) was added to the tube and nuclei were gently resuspended using a regular-bore P1000 pipette. The nuclei solution was then passed through a 40 uM strainer into a clean 2 mL LoBind tube to remove cell debris and larger clumps, then a 30 uM strainer into a clean 5 mL LoBind tube to further purify the sample. Nuclei were further purified using fluorescence-activated cell sorting to isolate nuclei based on size, shape, and DAPI+ fluorescence (a nuclear marker). Purified samples were immediately moved to library construction.

### Library Construction, Sequencing and Demultiplexing

Libraries were constructed and sequenced at the Petit Institute for Bioengineering and Bioscience at Georgia Tech. A cDNA library was constructed for each sample using the 10X Genomics Single Cell 3’ Chromium platform (v3.1; 10X Genomics, CA) (116). Libraries were sequenced on an Illumina NovaSeq using an S1 flow cell capturing 1.6 B paired-end ∼100bp reads, targeting ∼50k reads/nucleus. Supplementary Table S1 and Figures S1 and S2 summarize final sequencing parameters and demultiplexing results. The final mean read depth for each sample was ∼75k reads/nucleus, covering ∼1800 genes/nucleus. Sequencing saturation for all four samples ranged between 87.2-88.3%. Individual fish were matched to individual nuclei within each pooled sample using the demultiplexing algorithm souporcell (117). Souporcell identifies polymorphisms in nuclear RNAs and uses this information to predict groups of nuclei within a pool that came from the same individual animal.

### Quality control

FASTQ files were processed with Cell Ranger (10X Genomics) and reads were aligned to the *Amphiprion percula* genome assembly using a splice-aware alignment algorithm (STAR) within Cell Ranger. Gene annotations were obtained from the same assembly (ENSEMBL assembly accession: GCA_003047355.1, Nemo_v1). Because nuclear RNA contains introns, introns were included in the cellranger count step. Cell Ranger filtered out UMIs that were homopolymers, contained N, or contained any base with a quality score <10. Cell Ranger output included four filtered feature-barcode matrices (one per pool) containing expression data for a total of 24,480 features (annotated genes) and a total of 23,211 barcodes (corresponding to barcoded droplets containing nuclei) that were used passed through additional quality control steps in the ‘Seurat’ package in R. Total numbers of recovered transcripts and genes were highly similar in all four pools and so the same initial quality control filters were used across all pools. First, to minimize the impact of potentially dead or dying nuclei, barcodes associated with fewer than 600 total genes (n=1,216, ∼5%) were excluded. Next, to reduce the risk of doublets or multiplets, barcodes associated with more than 5,500 genes or 18,000 total transcripts (n=66, ∼0.2%) were also excluded. In total, 21,929 barcodes (94.5%) passed all quality control filters and were included in downstream clustering.

### Clustering and Cluster Annotation

Dimensionality reduction and clustering were performed in Seurat following a standard workflow. First, counts were log-normalized and the 4,000 most highly variable features were identified. Expression of these highly variable features were scaled (standardized as z-scores), and these scaled values were used for principal components analysis using the RunPCA function in Seurat. The first 50 principal components were then used for UMAP dimensionality reduction using the RunUMAP function in Seurat (minimum distance = 0.5, neighbors = 50). A shared nearest-neighbors graph was constructed from the first two UMAP dimensions (k = 50, no pruning). Clustering was then carried out at both low and high resolutions (0.02 and 1.3, respectively). Low-resolution clustering produced a set of 20 “parent” clusters, and high-resolution clustering produced a set of 49 “child” clusters nested within the parent clusters. Each parent cluster was assigned a number (1–20), and if a parent cluster contained multiple child clusters then each child cluster was assigned a letter (a, b, c, etc.). For example, parent cluster 14 contained four child clusters, designated 14a-d. Although olfactory bulbs were removed during dissection, we noticed a small parent cluster that bore strong gene expression signatures of olfactory bulb neurons (supported in part by SAMap alignment with cichlid snRNA-seq and spatial transcriptomics data, described below) and that was highly variable in relative proportions across individuals, consistent with variable and trace remnants of olfactory tissue being retained following olfactory bulb removal. Because the olfactory bulbs also contain radial glia, we independently reclustered radial glia and tested if any were strongly correlated in proportion with the proportion of putative olfactory neurons across individual subjects. Indeed, two small subclusters of radial glia whose proportions were strongly correlated with the proportion of this putative olfactory neuronal cluster across subjects. To minimize the influence of potential olfactory contamination in downstream analyses, these neuronal and radial glial nuclei (n=286 total) were excluded, yielding a total of 21,643 nuclei distributed across 19 parent and 48 child clusters that were included in final analyses.

### Cluster marker gene analysis

For each parent and child cluster, we identified genes that were differentially expressed among nuclei within the cluster compared to all other nuclei (hereafter, marker-DEGs) by performing a Wilcoxon Rank-Sum test using the FindMarkers function in Seurat (logfc.threshold=0, min.pct=0. To correct for multiple comparisons, we calculated q-values assuming a 5% false discovery rate using the ‘qvalue’ package in R, and genes were considered significant for a given cluster if q<0.05. Marker-DEGs were in turn used to assign clusters to major cell types (e.g. neurons, radial glia, etc.) and to identify which cell types expressed specific neurotransmitter, neuropeptide, or receptor genes of interest (see Supplementary Excel File 2 for a complete list of all *a priori* genes of interest).

### Cell-cell communication analysis

Cell-cell communication analysis was performed with the NeuronChat package for R. Briefly, NeuronChat uses a database of known protein-protein interactions that encompasses signaling ligands (and related vesicular transporters and synthesis enzymes) and receptors, as well as cell-cell adhesion complexes, and additionally tracks information on heterodimeric complexes and mediator proteins. It then analyzes gene expression between a sender cell population and a receiver cell population to estimate the directional (sender-to-receiver) potential for molecular communication between those populations. To align anemonefish genes with this database, we used their predicted mammalian homologs. Cell population labels included all child clusters except 2a and 13d. Child cluster 2a was split into the relatively mature and immature subpopulations, and cluster 13d was split into 47 subpopulations that were defined by *a priori* genes of interest (we excluded genes of interest that were not detected in all individuals within parent cluster 13). Because some nuclei within 13d co-expressed multiple genes of interest, these nuclei were represented across multiple subpopulations. Communication weights were estimated following a standard workflow with default parameters. Communication weights for each pairwise combination of cell populations were calculated using the createNeuronChat, run_NeuronChat, and net_aggregation functions with default parameters. Molecular systems underlying predicted communication patterns were further examined and visualized using NeuronChat as well as the ‘ggalluvial’ and ‘circlize’ packages in R.

### Neuroanatomical inference

To investigate the neuroanatomical origins of anemonefish snRNA-seq clusters, we first investigated expression patterns of known neuroanatomically-restricted *a priori* genes of interest (e.g. dorsal pallial marker genes, subpallial marker genes, hypothalamic and neuropeptide-related marker genes; see Supplementary Excel File 2). As a second and complementary line of investigation, we used SAMap (56), a Python tool that uses a self-assembling manifold (SAM) algorithm specifically designed to map cell atlases between species. SAMap weighs gene sequence similarities between species during integration of the data onto a shared, reduced-dimensional space, prioritizing genes with higher sequence similarity. Protein sequences were obtained from the same reference genome assemblies corresponding to the underlying snRNA-seq data (for anemonefish, Esembl genome assembly Nemo_v1, GCA_003047355.1; for cichlids, NCBI RefSeq assembly M_zebra_UMD2a, GCF_000238955.4). The map_genes.sh bash script from SAMap was used to perform blastp analysis, identifying reciprocal blast hits between anemonefish and cichlids. Raw gene count matrices from the underlying anemonefish and cichlid snRNA-seq datasets were each converted into .h5ad format and processed using the run function in SAMap with otherwise default settings. Cross-species mutual top hit cell type pairs were considered homologous. Anatomical locations were inferred based on neuroanatomical genetic markers described above as well as spatial mapping of the homologous cichlid cell population onto previously published cichlid spatial transcriptomics data (54). Inferences about the anatomical source of cell clusters further reinforced assignments of nuclei as olfactory-derived during quality control (see above in “Clustering and Cluster Annotation”).

### Sex Differences in Cluster-Specific Gene Expression

Sex differences in gene expression within clusters were analyzed using a negative binomial mixed regression model using the ‘glmer’ package in R. The outcome variable was the number of transcripts of a gene per nucleus. Sex was entered as a fixed effect and individual as a random effect. Dispersions were estimated using a Gamma-Poisson model. The natural logarithm of the library sizes per nucleus was entered as an offset term to normalize variation in sequencing depth across the nuclei. As above, the threshold for significance was based on a 5% FDR across all genes within clusters (q < 0.05).

Clusters were tested to determine whether they were overrepresented for sex-DEGs using a fisher exact test in R. Specifically, the proportion of sex-DEGs in a cluster (out of the total number of genes measured in the cluster) was compared to the proportion of sex-DEGs in all other clusters (out of all the other genes measured). We also tested whether there was a 50-50 split in the number of sex-DEGs that were upregulated and downregulated in females versus males, or whether there was a significant bias in the direction of the sex differences using a chi-square goodness of fit test. As above, the threshold for significance was q<0.05.

### Functional enrichment testing

Enrichment of biological functions among sex-DEGs was analyzed using the ClusterProfiler package in R. The enrichGO, enrichDO, and enrichKEGG functions were used to investigate overrepresentation of Gene Ontology (GO) terms, Human Diseases, and Kyoto Encyclopedia of Genes and Genomes (KEGG) pathways among FDR-significant sex-DEGs (q<0.05) in each parent and child cluster. Categories that were significantly overrepresented after accounting for a 5% false discovery rate were considered enriched (q<0.05). Differential enrichment was analyzed by cluster using a Fisher’s Exact Test. The test compared the number of upregulated sex-DEGs in males in a given category, the number of upregulated sex-DEGs in females in that category, the number of upregulated sex-DEGs in males not in that category, and the number of upregulated sex-DEGs in females not in that category. Categories were considered differentially enriched if they were significant after adjusting for a 5% false discovery rate (q<0.05).

### Identification of EREs, AREs, and GREs in the anemonefish genome

Estrogen, androgen, and glucocorticoid receptors function as hormone-responsive transcription factors that modulate gene expression by attaching to specific DNA segments known as estrogen, androgen, and glucocorticoid response elements, respectively. These elements are by characteristic nucleotide sequence motifs containing a three-base gap (ERE: AGGTCA---TGACCT; ARE: GGTACA---TGTTCT; GRE: AGAACA---TGTTCT). We identified genes located within a 25 kb radius of these canonical sequence motifs using the same reference genome. To determine the gene nearest to each response element, we employed the ’closest’ function in bedtools version 2.29.1. Genes closest to these elements were then annotated as ERE-, ARE-, and/or GRE-containing genes and used in downstream analyses.

### Enrichment of ERE, ARE, and GRE genes among sex-DEGs

To determine whether genes with EREs, AREs, and GREs were enriched among the sex-DEGs in each cluster, we used a Fisher’s Exact test in R (a separate test was conducted for each of the three response elements). For each cluster, the proportion of sex-DEGs containing the response element of interest (ERE, ARE, or GRE) was compared to the proportion of all other genes containing the response element of interest. Raw p-values were converted to q-values using the qvalue package in R, and the threshold for significance was q<0.05.

### Sex Differences in Cluster Proportions

For parent and child clusters, sex differences in cluster proportions were analyzed using a mixed effects binomial regression model using the ‘glmer’ package in R. In the model, the outcome variable was nuclei that either belonged to the cluster of interest or not. Sex was treated as a fixed effect, and the individual animal from which the nucleus was derived was treated as a random effect. Cluster proportions were also evaluated for specific neuropeptide and steroid receptor positive cells in POA cluster 13d. For these analyses, the same approach was used except that nuclei outside of cluster 13d were excluded. Thus, the outcome variable represented the total number of nuclei positive for the neuropeptide/steroid receptor gene out of the total number of nuclei in cluster 13d. Similarly, for proportion analyses of radial glial subclusters, the outcome variable represented the total number of nuclei belonging to each subcluster out of the total number of nuclei in parent cluster 14. For all analyses, sex differences in cluster proportions were considered significant if q < 0.05.

### Reclustering of neuronal and radial glial subpopulations

To investigate subpopulations within child cluster 2a, nuclei within this cluster were reclustered independently from all other nuclei in the dataset. Reclustering was performed using the following functions and parameters in Seurat: FindVariableFeatures (selection.method = “vst”, nfeatures = 2000), ScaleData, RunPCA (dim = 50), RunUMAP (dims = 1:50, min.dist = 0.5, n.neighbors = 50, metric = “euclidean”), FindNeighbors (reduction = “pca”, k.param = 50, dims = 1:2, prune.SNN = 0), and FindClusters (resolution = 0.2, algorithm = 2). A similar general approach was used to investigate subpopulations of radial glia. Nuclei within parent cluster 14 were reclustered independently from all other nuclei in the dataset using the following functions and parameters in Seurat: SCTransform, RunPCA (dim = 50), RunUMAP (dims = 1:10, min.dist = 0.5, n.neighbors = 50, metric = “euclidean”), FindNeighbors (reduction = “pca”, k.param = 50, dims = 1:10, prune.SNN = 0), and FindClusters (resolution = 1, algorithm = 2).

### Sex Differences in Predicted Cellular Maturity

To estimate predicted cellular maturity, we used the CytoTRACE package for R (118). Briefly, this tool uses transcriptional diversity as a relative indicator of cellular maturity, and assigns each nucleus a predicted maturity score ranging from 0 (relatively more mature) to 1 (relatively more immature). Sex differences in CytoTRACE score were analyzed using a linear mixed effects regression model in R, in which CytoTRACE score for each nucleus was the outcome variable, sex was a categorical fixed effect, and the individual from which each nucleus originated was considered a random effect. Effects were considered significant if q<0.05.

### Analysis of radial glial functional states

Differences in biological function across radial glial subclusters were investigated using CytoTRACE score as a measure of predicted cellular maturity (described above in “Sex Differences in Predicted Cellular Maturity”), as well as established genetic markers of radial glial quiescence, cycling, and commitment to neuronal differentiation (Supplementary Excel File 12). Differences were analyzed using a Kruskal Wallis test of each score (outcome) across subclusters (group), and a Dunn Test was used to identify significant pairwise differences among subclusters.

### Identification of IEG-like genes

To maximize power for tracking transcriptional signatures of neuronal excitation, we followed an approach outlined previously in Johnson et. al (42). Briefly, we identified a set of genes that were selectively co-expressed with each of three canonical IEGs (*egr1*, *npas4a*, and *fosab*) across many child neuronal clusters independently. For each of these three IEGs, nuclei within each cluster were split into IEG-positive versus IEG-negative. Genes that were differentially expressed between IEG-positive versus IEG-negative nuclei were identified using the FindMarkers function in Seurat, with “logfc.threshold” set to 0, and “min.pct” set to 1/54 (we used 54 because this was the number of nuclei in the smallest neuronal cluster). Within each cluster, any genes that did not meet this criterion were excluded and were assigned a *p* value of 1. Because FindMarkers requires at least three nuclei to be present in both comparison groups, clusters that contained less than three IEG-positive nuclei were excluded. Genes that were detected in the majority of clusters, and that were significantly upregulated (*p*lJ<lJ0.05) in IEG-positive nuclei in the majority of those clusters were considered to be significantly co-expressed with each individual IEG. Finally, only those genes that were significantly co-expressed with all three IEGs were considered to be IEG-like genes. This resulted in a set of 22 unique IEG-like genes that were used for downstream analyses (Supplementary Excel File 19).

### Sex Differences in IEG Expression

Sex-associated IEG expression was analyzed in parent clusters, child clusters, and neuropeptide/steroid hormone-related gene-defined populations within cluster 13d. Each nucleus was assigned an IEG score equal to the number of unique IEG-like genes (*n*lJ=lJ22) expressed. Sex differences in IEG score were analyzed in each cluster or population using a generalized linear mixed-effects regression model assuming a negative binomial distribution using the glmer.nb function for the lmer package in R. In the model, the outcome was the number of IEG-like genes observed in each nucleus, sex as well as the log of the total number of unique genes observed in the nucleus were treated as categorical and continuous fixed effects, respectively, and the individual from which each nucleus originated was treated as a random effect. Effects were considered significant if q<0.05.

### Sex differences in predicted cell-cell communication

To measure sex differences in predicted neuronal communication, we recalculated NeuronChat weights within each individual using the same approach described above except 1) non-neuronal populations were excluded and 2), we used M=200 in the run_NeuronChat step to reduce noise. We then scaled NeuronChat weights within each individual, and observed a bimodal-like distribution with one large peak near zero and a second peak near 1. To minimize the risk of potentially false-positive signals caused by baseline expression of compatible interaction genes between random neuronal populations, we excluded the >75% of population pairs with weights in the lower distribution by applying a cutoff of 0.65. For remaining pairwise combinations of cell population pairs (n=675), we analyzed sex differences in NeuronChat weights with a two-tailed permutation test using the twoSamplePermutationTestLocation function in the EnvStats package for R, with n.permutations = 10,000. Because a binomial test revealed a disproportionate number of results exhibiting a two-tailed p-value<0.05, population pairs passing this threshold were included in further downstream analyses of sex differences in cell-cell communication. To test for enrichment of hub populations among significant hits, we defined hub populations as neuronal populations that were identified as significant hits by multiple lines of earlier analyses, including analyses of sex-associated gene expression, sex-associated proportion differences, and sex-associated IEG expression. We then used a Fisher’s Exact Test to test if pairs exhibiting sex differences in predicted communication were enriched for connections in which a hub population was both a sender and receiver compared to pairs that did not exhibit sex differences in predicted communication (two-tailed p>0.05). Similarly, we used a Fisher’s Exact Test to identify populations that were overrepresented as senders and receivers among pairs exhibiting sex differences in predicted communication compared to pairs that did not exhibit sex differences in predicted communication.

## Supporting information

Supplementary Results

## Acknowledgments

This work was supported in part by NIH R01 GM144560 (JTS) and in part by OD P51OD011132 to Emory National Primate Research Center (ZVJ). Thanks to Peter Fenton for providing the funding for the single nuclei RNA sequencing, and for his passionate engagement in science, philosophy, and natural discovery.

