## Supplementary Results for "Adult sex change leads to extensive forebrain reorganization in clownfish"

Functional enrichment testing of sex-DEGs

Compared to other genes expressed in the forebrain, sex-DEGs were enriched for a large number of GO categories related to ribosomal structure and function as well as synaptic organization, plasticity, and signaling across many clusters. In specific clusters, sex-DEGs were enriched for GO terms such as “anchoring junction” (microglial cluster 17), “calcium-dependent protein binding” (radial glial cluster 14b), “ephrin receptor activity” (glutamatergic neuronal clusters 1a and 2a), “GABA-A receptor complex” (cluster 2a), and “chloride channel complex” (cluster 2a); as well as multiple categories related to endocrine and reproductive diseases (cluster 13b) (Supplementary Excel Files 4 and 5). Taken together, these results show widespread sex differences in cell type-specific gene expression spanning neurons, microglia, radial glia, and oligodendrocytes, and highlight cell type-specific specific molecular signaling pathways that may be functionally divergent between males and females.

Functional enrichment test of male versus female upregulated sex-DEGs

Enrichment testing revealed that sex-DEGs that were upregulated in females versus males (and vice versa) were related to specific biological functions (Supplementary Excel Files 4 and 5). For example, in parent clusters 1, 2, and 15, sex-DEGs that were upregulated in females were differentially enriched (compared to sex-DEGs that were upregulated in males) for multiple GO categories related to apoptosis (clusters 2 and 15) or autophagy (cluster 1), whereas in these same clusters, sex-DEGs that were upregulated in males were differentially enriched for multiple categories related to neuronal differentiation. Taken together, these results highlight specific biological functions and processes, e.g. neurogenesis and cell death, that may be involved in maintaining sex differences in forebrain cellular composition.

Response elements among sex-DEGs

We hypothesized that sex-associated gene expression across telencephalic cell populations may be regulated in part by sex differences in steroid hormones. Briefly, steroid hormones (e.g. estrogen, androgens, and glucocorticoids) are known to bind target receptors and form transcription factor complexes that enter the nucleus and bind directly to DNA sequence motifs (estrogen response elements, EREs; androgen response elements, AREs; and glucocorticoid response elements, GREs) to regulate transcription of nearby genes. Male *A. ocellaris* differ from females in levels of circulating sex steroid hormones such as estradiol (estrogen) and 11-ketotestosterone (the main bioactive androgen in fish), and display a trend for higher levels of glucocorticoids (57). To investigate the extent to which steroid hormones may explain sex-associated gene expression, we identified genes containing predicted EREs, AREs, and GREs and tested if they were overrepresented among sex-DEGs across clusters. Compared to other genes in the genome, sex-DEGs were significantly more likely to reside near steroid response elements in parent clusters 1 (AREs, GREs), 2 (GREs), 7 (EREs), 9 (EREs, GREs), and 15 (EREs, AREs, GREs) (Supplementary Excel File 6. These patterns were largely mirrored among child clusters: sex-DEGs were enriched for steroid response elements in glutamatergic clusters 1a (EREs) and 2a (EREs); GABAergic cluster 9a (EREs, GREs); radial glial cluster 14b (GREs); and oligodendrocyte cluster 15a (AREs) (Supplementary Excel File 7). However, this pattern only captured a minority of overall sex-DEG effects and clusters, suggesting that other mechanisms are likely important in regulating sex-associated gene expression. Nonetheless, these results are consistent with specific steroid hormones contributing to sex-associated gene expression in specific populations of neurons and glia throughout the forebrain, and with previous work demonstrating a functional role for sex steroid hormones in sex change (33, 58, 59).
